## Supplementary material for "Reliable RNA-seq analysis from FFPE specimens as a means to accelerate cancer-related health disparities research": Revised Supplemental Methods

### Supplementary Methods

RNA and DNA extraction from formalin-fixed paraffin-embedded (FFPE) tissue samples can be a challenging process due to the cross-linking and degradation of RNA/DNA caused by formalin fixation. However, we demonstrated with the right protocol, it is possible to extract RNA with sufficient quality to generate reliable sequencing data as well as residual DNA recovered during RNA isolation with adequate quality for SNP array platforms.

#### 1. RNA extraction from formalin-fixed paraffin-embedded (FFPE) tissue

We used AllPrep DNA/RNA FFPE, commercially available kit, from Qiagen (Catalog No. 80234) and some modifications to the manufacture instructions were made.

Sample Preparation:

- FFPE slide sections were cut to 6 µm thickness on glass slides, prepared from archival blocks, containing oropharyngeal squamous cell carcinomas (OPSCC) specimens, which ranged in storage time from 1 to 20 years.
- Alternatively, sections could be sliced directly into Eppendorf tubes and stored at room temperature in a desiccator prior to processing.
- Parameters to consider: average area of tissue per slide= 2 sq.cm, average number of slides simultaneously processed= ~9, average total area of tissue processed= 18 sq.cm, and average yield per total area of tissue= 402 ng/sq.cm (median= 274 ng/sq. cm).
- In case > 10 slides are needed, then cut more slides or proceed with a punch biopsy.
- Trim excess of paraffin off the sample block as much as possible.
- Use a fresh flat edge razor to scrape tissue sections from the slides into a sterile microcentrifuge tube.
- To remove the paraffin penetrated into the tissue, add 1 mL of xylene and vigorously vortex for 10 seconds.
- Centrifuge at 14,000 x g for 2 minutes at room temperature and carefully remove the xylene without disturbing the pellet.
- Wash the tissue pellet twice with 100% ethanol (1mL). Re-pellet during each wash by centrifugation at 14,000 x g for 2 minutes at room temperature, and discarding supernatants. Remove residual xylene after second wash and air-dry the pellet for 5-10 minutes.

Tissue Digestion:

- Resuspend the pellet by adding 150 µL of buffer PKD and flicking the tube to loosen the pellet.
- Add 10 µL of proteinase K and mix by vortexing.
- Incubate at 56°C for 15 minutes on a heating block.
- Incubate on ice for 3 minutes.
- Centrifuge at 14,000 x g for 15 minutes.
- Carefully transfer the supernatant, without disturbing the pellet, to a new 2.0 mL safe-lock microcentrifuge tube

Note: The supernatant contains RNA and the pellet contains DNA.

- Store the pellet at -20°C for subsequent DNA purification.

Purification of total RNA:

- Incubate the supernatant from the previous step at 80°C for 15 minutes on a heating block.
- Briefly centrifuge the tube to remove drops from the inside of the lid.
- Add 320 µL of Buffer RLT to adjust binding conditions, and mix by vortexing,
- Add 1120 µL 100% ethanol and mix well by vortexing for 5 seconds.
- Transfer 700 µL of the sample to an RNeasy MinElute spin column placed in a 2 mL collection tube. Close the lid gently.
- Centrifuge at 8000 x g for 15 seconds and discard the flow-through from the collection tube. Reuse the collection tube in the next step.
- Repeat the previous step until the entire sample has passed through the RNeasy MinElute spin column. Reuse the collection tube in the next step.
- Add 350 µL Buffer FRN to the RNeasy MinElute spin column. Close the lid gently.

  Note: Before using Buffer FRN for the first time, check whether a precipitate has formed. If necessary, dissolve by warming with gentle agitation. After equilibration to room temperature (15–25°C), add 42 ml isopropanol (96–100%) to the entire concentrate (14 mL). Label the bottle.
- Centrifuge at 8000 x g for 15 seconds. Discard the flow-through. Place the RNeasy MinElute spin column in a new collection tube.
- Add 80 µL DNase I solution directly to the RNeasy MinElute spin column membrane, and place on the benchtop (20–30°C) for 15 min.

  Note: 1- Prepare DNase I stock solution by dissolving the lyophilized DNase I (1500 Kunitz units) in 550 µL RNase- free water. In some cases, the vial of DNase may appear to be empty. This is due to lyophilized enzyme sticking to the septum. To avoid loss of DNase I do not open the vial. Inject RNase-free water into the vial using an RNase-free needle and syringe. Mix gently by inverting the vial. Do not vortex.

  2- Prepare DNase I solution by adding 10 µL of DNase I stock solution into 70uL of Buffer RDD.

  3- DNase I is especially sensitive to physical denaturation. Mixing should only be carried out by gently inverting the tube. Do not vortex.
- Add 500 µl Buffer FRN to the RNeasy MinElute spin column. Close the lid gently.

  Note: Before using Buffer FRN for the first time, check whether a precipitate has formed. If necessary, dissolve by warming with gentle agitation. After equilibration to room temperature (15–25°C), add 42 mL isopropanol (96–100%) to the entire concentrate (14 mL). Label the bottle.
- Centrifuge at 8000 x g for 15 seconds. Save the flow-through for use in the next step.

  Note: Do not discard the flow-through, as it contains RNA including small RNAs.

- Place the same RNeasy MinElute spin column in a new 2 mL collection tube. Apply the flow-through from the previous step to the RNeasy MinElute spin column. Close the lid gently.
- Centrifuge at 8000 x g 15 seconds. Discard the flow-through. Reuse the collection tube in the next step.
  Note: RNA is bound to the RNeasy MinElute spin column membrane.
- Add 500 µL Buffer RPE to the RNeasy MinElute spin column. Close the lid gently.
  Note: Add 4 volumes (44 mL) ethanol (100% and molecular biology grade) to the bottle containing 11 mL Buffer RPE concentrate. Label the bottle. Before starting the procedure, mix reconstituted Buffer RPE by shaking.
- Centrifuge at 8000 x g for 15 seconds to wash the spin column membrane. Discard the flow-through. Reuse the collection tube in the next step.
- Add 500 µL Buffer RPE to the RNeasy MinElute spin column. Close the lid gently.
- Centrifuge at 8000 x g for 15 seconds to wash the spin column membrane. Discard the flow-through.
- Place the RNeasy MinElute spin column in the same collection tube.
- Centrifuge at 10,000 x g for 2 minutes. Discard the collection tube with the flow-through.
- Place the RNeasy MinElute spin column in a new 1.5 mL collection tube.
- Add 15–30 µL RNase-free water directly to the RNeasy MinElute spin column membrane.
- Incubate for 2 minutes at room temperature.
- Centrifuge at 10000 x g for 1 minute to elute the RNA.
  Note: RNA is now in the flow- through, do not discard.
- Preserve the DNA pellet at -20°C.

RNA Quality Control:

- Measure the concentration and assess the quality of the extracted RNA using Nanodrop (Cytation 3 imaging reader and Gen5 Image 2.07 software).
  Note: ratio of 260/280 should be 1.8-2.0 and ration of 260/230 is preferred to be 2.0-2.2.

### 2. Purification of genomic DNA

We used COBAS DNA Sample Preparation Kit, commercially available, from Roche (Catalog No. 05985536190) and some modifications to the manufacture instructions were made.

- The DNA pellet preserved at -20°C should be equilibrated first to room temperature prior to start the DNA purification protocol.
- Resuspend the DNA pellet in 180 µL DNA TLB and mix for 10 seconds by vortexing.
- Add 70 µL of reconstituted Proteinase K to the same 1.5 mL microcentrifuge tube and vortex for 10 seconds.
- If required for the test, place a negative control tube adding 180 µL DNA TLB and 70 µL of reconstituted Proteinase K. Vortex for 10 seconds.

Note: The negative control should be processed following the same procedure as the samples.

- Place tubes in 56°C incubator with agitation/rotation for 1 hr.
- Place tubes in the 90°C dry heat block and incubate for 1h.
- Allow the tubes to cool to 15°C to 30°C.
- Briefly centrifuge the tubes to remove drops from the inside of the lid.
- Add 200 µL DNA PBB to each tube, mix by pipetting and incubate the tubes at room temperature for 10 minutes.
- Add 100 µL isopropanol to each tube and mix lysate by pipetting.
- Transfer the lysate into a Filter Tube (FT)/Collection Tube (CT) unit.
- Centrifuge the FT/CT units at 8,000 x g for 1 minute.
- Place the FT in a new CT and discard the old CT with the flow-through.
- Add 500 µL working WB I to each FT.
  Note: Prepare working WB I by adding 15 mL of absolute ethanol to the bottle of WB I. Mix by inverting the bottle 5 to 10 times. Label de bottle. Store working WB I at 15 to 30 for up to 90 days or until the expiration date, whichever comes first.
- Centrifuge FT/CT units at 8,000 x g for 1 minute. Discard the flow-through. Place the FT back into the same CT.
- Add 500 µL working WB II to each FT.
  Note: Prepare working WB II by adding 50 mL of absolute ethanol to the bottle of WB II. Mix by inverting the bottle 5 to 10 times. Label de bottle. Store working WB I at 15 to 30 for up to 90 days or until the expiration date, whichever comes first.
- Centrifuge FT/CT units at 8,000 x g for 1 minute. Discard the flow-through. Place the FT into a new CT.
- Centrifuge FT/CT units at 14,000 x g for 1 minute to dry the filter membranes. Discard the CT with the flow-through.
- Place each FT into an elution tube (1.5 mL microcentrifuge tube).
- Add 100 µL DNA EB directly to the FT membrane.
- Incubate the FT with elution tube 15°C to 30°C for 5 minutes.
- Centrifuge the FT with elution tube at 8,000 x g for 1 minute to collect eluate into the elution tube.
  Note: Eluted DNA is now in the flow-through, do not discard.
- Preserve the elution tube containing the DNA stock at -20°C until quantification.

<u>DNA quantification</u> (First PicoGreen):

We used Quant-iT PicoGreen dsDNA Reagent and Kit, commercially available, from Invitrogen (Catalog No. P11496) and some modifications to the manufacture instructions were made.

- Prepare the DNA standard curve from DNA standard original stock 100 µg/mL storage at 4°C.
- Prepare 8 tubes and label A to H for the standards.

- Prepare 20 mL of 1x working solution of TE from 20x TE. Use the nuclease free water for the dilution.
- See below appropriate volume of TE for each standard dilutions.
  Add 495 µL of TE to tube A.
  Add 250 µL of TE to tubes B, D and E
  Add 400 µL of TE to tube C
  Add 450 µL of TE to tubes F to H
- Perform dilution of standard as follows:
  Add 5 µL of Standard to tube A, mix by pipetting, and take 250 µL to transfer this volume to tube B.

  Note: Change tips between dilutions. The dilutions must be mixed vigorously before transferring to the next tube for serial dilution.

  Take 100 µL from tube B and transfer the volume to tube C
  Take 250 µL from tube C and transfer the volume to tube D
  Take 250 µL from tube D and transfer the volume to tube E
  Take 50 µL from tube E and transfer the volume to tube F
  Take 50 µL from tube F and transfer the volume to tube G
- Final concentrations of the standards is as follows:

| | | **Previous Tube volume (µL)** | **Serial Dilution** | **Concentration (ng/mL)** |
|---|---|---|---|---|
| **Tube** | **1X TE (µL)** | | **Dilution factor** | 100000 |
| A | 495 | 5 | 100 | 1000 |
| B | 250 | 250 | 2 | 500 |
| C | 400 | 100 | 5 | 100 |
| D | 250 | 250 | 2 | 50 |
| E | 250 | 250 | 2 | 25 |
| F | 450 | 50 | 10 | 2.5 |
| G | 450 | 50 | 10 | 0.25 |
| H | 450 | Blanck (negative control) | | |

- Tube H will only have TE. This is the blank control.
- Picogreen comes in a 100 µL stock volume. Prepare a working solution with 1:200 ratio. Add 50 µL of picogreen solution into 9.95 mL of 1x TE.
- Using a multi-channel pipet, add 100 µL of picogreen solution to each well in a 96 well plate and cover with aluminum foil.
- Draw a plate schematic making duplicates of standard and triplicate of sample to be quantified.
- For DNA samples to be quantified perform a 1:50 dilution ratio with 10 µL DNA sample plus 490 µL 1x TE.
- Mix very well and add 100 µL per well in the PicoGreen assay.
- Using Cytation 3 imaging reader and Gen5 Image 2.07 software, read the plate using a fluorescence at 480 excitation and 520 emission.

Read speed set to “normal” with 100 msec delay between reads. Set 2 gains: one automatic and second read set to 100 gain. With read height of 7mm.

- After read the plate, plot standard curve, and interpolate the samples to obtain protein measurement.
- Return the DNA samples to -20°C after quantification.
  Note: if the total preliminary DNA yield for a sample at this point is <350 ng, then process additional slides. The yields can be combined before the final DNA precipitation step. The goal is to achieve a minimal of 300 ng total DNA in the end of the process.

<u>DNA precipitation</u>:

- Thaw DNA sample on water if previously frozen.
  Note: If using two samples from the same patient because of insufficient yield, then pool the two samples before starting the ethanol precipitation procedure and make a note that volumes below need to be scaled up 2-fold or the doubled sample volume.
- Add 1/10 DNA sample volume of 3 M sodium acetate to DNA solution (e.g., 9 µL NaAC per 90 µL remaining DNA sample after quantification).
- Add 2 µL linear-acrylamide (stock 5mg/mL) to DNA/sodium acetate mixture.
- Add 2.5 volume of ice cold (-20°C) 100% ethanol (not isopropyl) to the mixture of DNA solution/sodium acetate/linear-acrylamide. [2.5 vol. x (vol. DNA solution + vol. sodium acetate + vol. linear-acrylamide)].
- Mix gently but thoroughly, inverting 20 times.
- Incubate the mixture at -20°C for 1 hr.
- Centrifuge the mixture at 14,000 x g for 15 minutes in 4°C centrifuge.
- Wash the DNA pellet by adding room temperature 70% ethanol made from nuclease free water. Close cap and flick tube slightly to expose bottom of pellet to ethanol wash.
- Centrifuge the mixture at 14,000 x g for 15 minutes in a room temperature centrifuge to keep salt dissolved.
- Leave the tube upside-down opened for 5 minutes or as needed to facilitate drying. Be careful with the position of the pellet on the tube. Pellets can become nearly invisible after drying.
- Dissolve DNA in 10 mM nuclease free Tris, pH 8.0.
  Note: 1-When deciding on final volume to solubilize DNA, calculate what would be expected to produce 100 ng/µL based on total yield from first PicoGreen.
  2- If DNA is already between 50 ng-100 ng/µL (result from the first PicoGreen) not dilute the sample.
  3- Otherwise, if the stock DNA is above 100 ng/µL, then proceed with the dilute down to 100ng/µL with 10 mM nuclease free Tris pH, 8.0.

<u>DNA quantification after precipitation and concentration</u>: (Second PicoGreen)

We used Quant-iT PicoGreen dsDNA Reagent and Kit, commercially available, from Invitrogen (Catalog No. P11496) and some modifications to the manufacture instructions were made.

- Perform the same procedure than in First PicoGreen

- For DNA samples to be quantified make a 1:500 dilution ratio with 1µL DNA sample plus 490 µL 1x TE.
- Mix very well and add 100 µL per well in the PicoGreen assay.

Sample DNA yields before ethanol precipitation

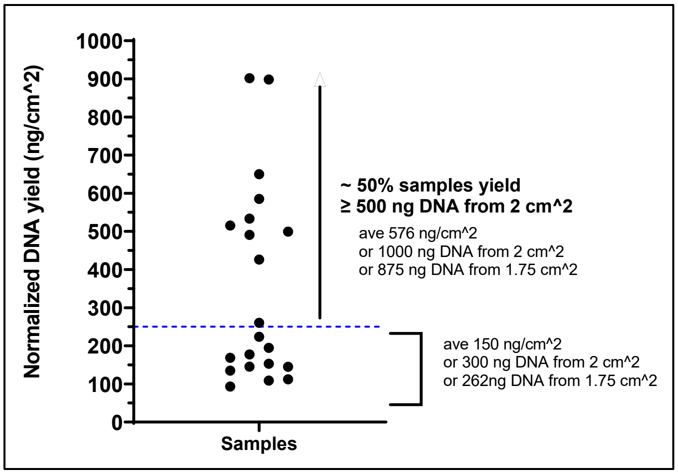


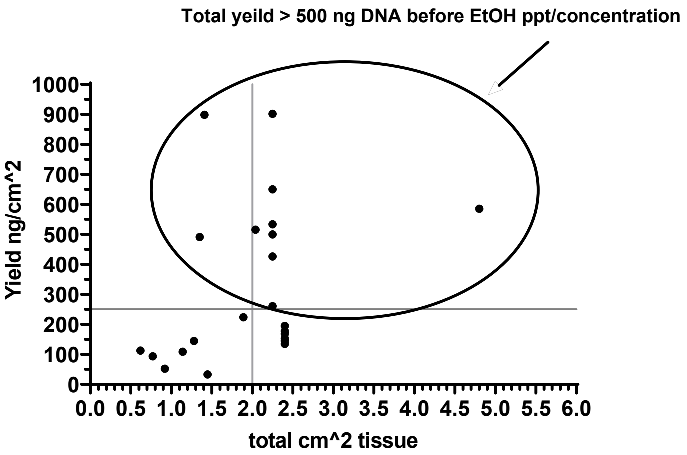


### 3. Codes utilized in analysis

**Overview of code utilized in analysis:** Four code packages are used in the pipeline for the following tasks:

**Requirements:** Install integrated development environment or interpreter for Python. We use both JupyterLab (.ipynb files) and Visual Basics (.py) as explained below, with dependencies import numpy, import pandas, import scipy.stats, and import binom libraries.

1) **Convert_tab.ipynb-** extracts and merges raw count data and effective gene sizes from individual *tab delimited* sample files within the "inputs" folder and concatenates data into a dataframe metafile file that can subsequently be used as input for the normalization program.

2) **Convert_csv.ipynb**- extracts and merges raw count data and effective gene sizes from individual *csv (comma delimited)* sample files within the "inputs" folder and concatenates data into a dataframe metafile file that can subsequently be used as input for the normalization program. The two Convert$.ipynb files are redundant and differ only by the type of input files they require.

3) **Normalize.py**- takes the csv outputs from the Convert$ programs and performs all the steps for the normalization pipeline, including FPKM-UQ normalizations, identification of technical outliers, global adjustment of cohort median, replacement of negative log2 values with zeros, outputs final log2 adjusted FPKM values, and linear FPKM values.

4) **Dropout.ipynb**-takes inputs of observed number of samples with zero expression for all genes from the study cohort and estimated or modeled probabilities of obtaining a zero value based on historical prevalence or reference, and determines the binomial probability and adjusted probability of obtaining at least the same number of observed zeros.

**Running Convert$.ipynb- Run with Jupyter notebooks**

1. Create a folder called "Converter" and subfolder within called "inputs".
2. Ensure raw count files include headers "gene_id", "effective_length", and "expected_count", with careful attention to matching text format (e.g, no capitilzaitons). When using raw counts from HTSeq, append files with a new column to assign gene sizes based on reference genome and label "effective_length" as header.
3. Create a file named "File_ID_Sample_ID_map.csv" with two column headers "File id" and "Sample ID". In the File id column list all raw count file input names truncated before the very first period. For example, a file called "TA_2.genes.results" would be listed as "TA_2". In the second column under Sample ID, type a corresponding custom sample nickname to identify the sample such as "Treatment_1" or any desired alphanumeric string. This creates a map so the output dataframe metafile will have headers with the assigned sample names instead of the file names.
4. Create a reference file called "genesymbol.csv" with header "gene_id" which lists all **protein coding** gene Id's using the same gene identification system found in the raw input files. This could be an Entrez id, or an Ensembl ID for example. See supplementary table 4 for a list of all protein coding human genes with their Ensembl ID, Entrez ID, or official symbols. This reference file
4. Save the "File_ID_Sample_ID_map.csv" and "genesybmol.csv" files in the Converter folder.
5. Upload all the raw count files into the "inputs" subfolder
6. Identify the last suffix of raw input files, such as "results" from our example "TA_2.genes.results", which is the last string of text after the last ".". If files do not end with ".results" than modify the code line in the Convert$.ipynb file that reads "if filename. Endswith ('.results') and replace with appropriate file extension.
7. Open JupyterLab and load appropriate Convert$.ipynb to match either tab or csv delimited inputs and hit run.
8. The csv merged metafile output appears in a file named "final_dataframe.csv" that will have the first column with all gene id's, then will repeat for every sample with a column repeating the designated sample nickname, a column with the gene effective length, and a column with the expected count.
9. Rename the "final_dataframe.csv" file as "Combined_data.csv" which will be used in the normalization program.

**Running Normalization pipeline: Run with Visual basics**

1. Create a folder called "Normalization" and save the "Normalization.py" file.
2. Save the "Combined_data.csv" file from above in same folder
3. Create a file called "gene_reference.csv" with two column headers "gene_id", and "Symbol" observing strict punctuation. Under the "gene_id" column list all the same gene id's from the "genesymbol.csv"

used in the converter pipeline and in the second column labeled “Symbol” list the corresponding official gene symbols (See Supplementary table 4).
4. Run the Normalization.py program which will automatically look for the metafile input “Combined_data.csv” and “gene_reference.csv” files.
5. The normalization program will output multiple intermediate files that are self-explanatory by name. The sample nicknames are preserved in the file named “step15_final_adjusted_log2_fmpkm_uq.csv” as columns with headers “Sample_ID#”, where # is the sample index according to the order from the “Combined_data” input file beginning with an index = 0 through N-1. This can be used to map sample indexes back to the sample nicknames in subsequent files that utilize only the sample indexes. Among other useful files are : (1) step8_outliers.csv which will list the average MAD value for each sample and whether the sample is a technical outlier based on the adjusted P value with an FDR =0.01, which can be adjusted in the code if another threshold is desired; (2) Final_adjusted_log2_fpkm_uq.csv = log2 of final upper quartile adjusted normalized counts, with cohort global median set = 7, and negative log2 values replaced by zeros for consistency. The transformation Y = log2 (X=0.01) was used to derive the log values and all negative log values were replaced with “0”. These normalized values can be used for downstream analyses such as differential analysis of gene expression, PCA analysis, Combat, etc; 3) step18_log2_to_linear.csv= normalized values converted back to linear space but preserving zero values that replaced negative log2 values.

**Running the dropout program. Run in Jupyter notebooks**

1) Create a folder called “zeros”
2) Save file “zeros_binomial.ipynb” in the zeros folder
3) Create a csv input file named “zeros.csv” with headers “Symbol”, “Observed_zeros”, “Total_samples”, and “Probability” which for each gene will contain: the official gene symbol, the numerical integer of samples in the test cohort that have zero raw counts for a gene (calculated using the merged “final_dataframe.csv” file or “Combined_data.csv” and not the processed normalized data where negative log2 values are replaced), the total samples in the test cohort, and the fractional probability or prevalence of samples with zero gene expression either modeled or taken from a reference cohort like the TCGA. Note, if the observed_zero value is actually “0” because no test cohort sample had a zero, automatically replace it with a “1” so that it is possible for the program to calculate a cumulative binomial probability. The output file will be called Zeros_with_P_values.csv and will have the same first headers and row information as the input file but will be appended with a “Raw_P_value” column and a “Adjusted_P_value” column that used the Benjamini Hochberg correction for multiple testing with an FDR=0.1.

**Converter_tab**

```
import os
import pandas as pd

# Path to the folder containing your files
folder_path = os.path.join(os.getcwd(), 'inputs')
```

```
# Load the gene symbol file (assuming this file contains 'gene_id')
gene_symbol_df = pd.read_csv('genesymbol.csv')

# Load the gene ID to sample ID mapping file
mapping_df = pd.read_csv('File_ID_Sample_ID_map.csv')

# Create a dictionary from the mapping for faster lookups
file_id_to_sample_id = dict(zip(mapping_df['File Id'], mapping_df['Sample ID']))

# Initialize a list to hold the dataframes, starting with gene_symbol_df (which
will hold the 'gene_id' column)
dataframes = [gene_symbol_df]

# Iterate over each file in the folder
for filename in os.listdir(folder_path):
    if filename.endswith('.results') and 'genesymbol' not in filename and
'File_ID_Sample_ID_map' not in filename:
        # Extract the file ID prefix (before the first period)
        file_id_prefix = filename.split('.')[0]

        # Check if this file ID prefix is in the mapping dictionary
        if file_id_prefix in file_id_to_sample_id:
            # Load the tab-delimited .results file
            file_df = pd.read_csv(os.path.join(folder_path, filename),
delimiter='\t')  # Tab-delimited

            # Select only the 'gene_id', 'effective_length', and 'expected_count'
columns by name
            try:
                selected_columns = file_df[['gene_id', 'effective_length',
'expected_count']].copy()
            except KeyError as e:
                print(f"Error: One of the expected columns is missing in
{filename}: {e}")
                continue

            # Merge the .results file with the genesymbol file to only keep
matching gene_ids
            selected_columns = pd.merge(gene_symbol_df[['gene_id']],
selected_columns, on='gene_id', how='inner')

            # Add the Sample ID to the DataFrame based on the filename (mapping
file_id_prefix to Sample ID)
            sample_id = file_id_to_sample_id[file_id_prefix]
```

```
            selected_columns.insert(1, 'Sample_ID', sample_id)  # Insert
Sample_ID right after gene_id

            # Drop the 'gene_id' column from the results file (we already have it
in the gene_symbol_df)
            selected_columns = selected_columns.drop(columns=['gene_id'])

            # Append the dataframe to the list (without the redundant 'gene_id')
            dataframes.append(selected_columns)

# Concatenate all dataframes horizontally, preserving only the first 'gene_id'
column
final_df = pd.concat(dataframes, axis=1)

# Save the final dataframe to a CSV file
final_df.to_csv('final_dataframe.csv', index=False)
```

**Converter_csv**

```
import os
import pandas as pd

# Path to the folder containing your files
folder_path = os.path.join(os.getcwd(), 'inputs')

# Load the gene symbol file (assuming this file contains 'gene_id')
gene_symbol_df = pd.read_csv('genesymbol.csv')

# Load the gene ID to sample ID mapping file
mapping_df = pd.read_csv('File_ID_Sample_ID_map.csv')

# Create a dictionary from the mapping for faster lookups
file_id_to_sample_id = dict(zip(mapping_df['File Id'], mapping_df['Sample ID']))

# Initialize a list to hold the dataframes, starting with gene_symbol_df (which
will hold the 'gene_id' column)
dataframes = [gene_symbol_df]

# Iterate over each file in the folder
for filename in os.listdir(folder_path):
    if filename.endswith('.results') and 'genesymbol' not in filename and
'File_ID_Sample_ID_map' not in filename:
        # Extract the file ID prefix (before the first period)
        file_id_prefix = filename.split('.')[0]
```

```
        # Check if this file ID prefix is in the mapping dictionary
        if file_id_prefix in file_id_to_sample_id:
            # Load the CSV .results file (reading as a CSV file)
            file_df = pd.read_csv(os.path.join(folder_path, filename))  # Reading
as CSV

            # Select only the 'gene_id', 'effective_length', and 'expected_count'
columns by name
            try:
                selected_columns = file_df[['gene_id', 'effective_length',
'expected_count']].copy()
            except KeyError as e:
                print(f"Error: One of the expected columns is missing in
{filename}: {e}")
                continue

            # Merge the .results file with the genesymbol file to only keep
matching gene_ids
            selected_columns = pd.merge(gene_symbol_df[['gene_id']],
selected_columns, on='gene_id', how='inner')

            # Add the Sample ID to the DataFrame based on the filename (mapping
file_id_prefix to Sample ID)
            sample_id = file_id_to_sample_id[file_id_prefix]
            selected_columns.insert(1, 'Sample_ID', sample_id)  # Insert
Sample_ID right after gene_id

            # Drop the 'gene_id' column from the results file (we already have it
in the gene_symbol_df)
            selected_columns = selected_columns.drop(columns=['gene_id'])

            # Append the dataframe to the list (without the redundant 'gene_id')
            dataframes.append(selected_columns)

# Concatenate all dataframes horizontally, preserving only the first 'gene_id'
column
final_df = pd.concat(dataframes, axis=1)

# Save the final dataframe to a CSV file
final_df.to_csv('final_dataframe.csv', index=False)
```

**Normalize.py**

```
import os
```

```
import numpy as np
import pandas as pd
from scipy.stats import norm
from statsmodels.robust.norms import HuberT
from statsmodels.robust.robust_linear_model import RLM
from statsmodels.stats.multitest import multipletests


# Load the combined RNA-seq data
# Load the combined RNA-seq data and save as Pickle for faster access
def load_combined_data(file_path, pickle_file="combined_data.pkl"):
    if os.path.exists(pickle_file):
        print(f"[LOG] Loading combined RNA-seq data from Pickle file:
{pickle_file}")
        data = pd.read_pickle(pickle_file)
    else:
        print(f"[LOG] Loading combined RNA-seq data from CSV file: {file_path}")
        data = pd.read_csv(file_path)
        print(f"[LOG] Saving combined RNA-seq data as Pickle: {pickle_file}")
        data.to_pickle(pickle_file)  # Save the dataframe as Pickle for faster
reloading
    return data


# Load the gene reference file
def load_gene_reference(file_path):
    gene_reference = pd.read_csv(file_path)
    print(f"[LOG] Loaded gene reference data from {file_path}")
    return gene_reference


# Step 1: Filter out non-protein-coding genes and create output with additional
"count" column
def filter_protein_coding_genes(data, gene_reference):
    # Merge data with the gene reference to filter out non-protein-coding genes
    filtered_data = pd.merge(data, gene_reference, how="inner", on="gene_id")
    print("[LOG] Filtered data to include only protein-coding genes")

    # Add 'count' column as the first column (for demonstration, using row
numbers as counts)
    filtered_data.insert(0, "count", range(1, len(filtered_data) + 1))

    # Save the step 1 output
    filtered_data.to_csv("step1.csv", index=False)
    print("[LOG] Saved step 1 output to 'step1.csv'")
```

```
    return filtered_data


# Step 2: Calculate the 75th percentile of counts for each sample and create
step2_MUQ.csv
def calculate_75th_percentile(filtered_data):
    percentiles = {}

    # Identify expected_count columns (every third column starting from 4th
column)
    sample_columns = filtered_data.columns[4::3]

    for sample in sample_columns:
        # Ensure that the column data is numeric
        sample_data = pd.to_numeric(filtered_data[sample], errors="coerce")
        sample_data = sample_data.replace(0, np.nan)  # Exclude zero counts

        # Check if the column has valid numeric data
        if sample_data.notna().sum() > 0:
            s_75 = sample_data.quantile(0.75)
            percentiles[sample] = s_75
            print(f"[LOG] Calculated 75th percentile for {sample}: {s_75}")
        else:
            percentiles[sample] = np.nan
            print(f"[LOG] No valid data for {sample}, setting 75th percentile as
NaN.")

    # Calculate the Median Upper Quartile (MUQ)
    valid_percentiles = [v for v in percentiles.values() if not np.isnan(v)]
    muq = np.median(valid_percentiles) if valid_percentiles else np.nan

    # Extract sample names (corresponding to the samples like "SP17-2267-FSA10")
    sample_names = [
        filtered_data.columns[i - 2] for i in range(4,
len(filtered_data.columns), 3)
    ]

    # Create the step2_MUQ.csv output
    step2_data = pd.DataFrame(
        {
            "Sample": sample_names,
            "Count 75th PCTLS": [percentiles[sample] for sample in
sample_columns],
            "MUQ": [muq] + [""] * (len(sample_columns) - 1),
```

```
        }
    )
    step2_data.to_csv("step2_MUQ.csv", index=False)
    print("[LOG] Saved step 2 output to 'step2_MUQ.csv'")

    return percentiles, muq


# Step 3: Apply gene size transformation (max(x, 252)) and calculate the median
gene sizes
def apply_gene_size_transformation(data):
    # Identify effective_length columns (every third column starting from the
third column)
    length_columns = data.columns[3::3]

    # Apply max(effective_length, 252) to each effective length column
    for length_column in length_columns:
        data[length_column] = data[length_column].apply(lambda x: max(x, 252))
        print(f"[LOG] Applied max(effective_length, 252) to {length_column}")

    # Collect all transformed gene sizes
    all_lengths = []
    for length_column in length_columns:
        all_lengths.extend(data[length_column].tolist())

    overall_median = np.median(all_lengths)

    # Save the transformed data to step3_MGS.csv
    data.to_csv("step3_MGS.csv", index=False)
    print("[LOG] Saved step 3 output to 'step3_MGS.csv'")
    print(f"[LOG] Global Median Gene Size (MGS): {overall_median}")

    return data, overall_median


# Step 4: Normalize FPKM-UQ using vectorized operations
def normalize_fpkm_uq(
    data, percentiles, muq, global_mgs, file_name="step4_fpkm_uq.csv"
):
    normalized_data = data.copy()
    sample_columns = normalized_data.columns[4::3]  # Sample columns

    # Vectorized normalization: (reads / (gene_length * s75th)) * muq *
global_mgs
    for i, sample in enumerate(sample_columns):
```

```
        s75th = percentiles[sample]
        gene_lengths = normalized_data.iloc[:, 3::3].values[
            :, i
        ]  # Vector of gene lengths
        reads = normalized_data[sample].values

        # Vectorized calculation of normalized values
        normalized_values = (reads / (gene_lengths * s75th)) * muq * global_mgs
        normalized_data[sample] = normalized_values

    normalized_data.to_csv(file_name, index=False)
    print(f"[LOG] Saved normalized FPKM-UQ data to '{file_name}'")
    return normalized_data


# Step 5: Log2 transformation of FPKM-UQ
# Step 5: Log2 transformation using vectorized operations
def log2_transform_fpkm_uq(data, file_name="step5_log2_fpkm_uq.csv"):
    log2_data = data.copy()
    fpkm_uq_columns = log2_data.columns[4::3]  # Sample columns

    # Vectorized log2 transformation for all sample columns at once
    log2_data[fpkm_uq_columns] = np.log2(log2_data[fpkm_uq_columns] + 0.01)

    log2_data.to_csv(file_name, index=False)
    print(f"[LOG] Saved log2 transformed FPKM-UQ data to '{file_name}'")
    return log2_data


# Step 6: Calculate MAD and identify outliers
def calculate_mad_and_identify_outliers(data,
file_name="step6_mad_outliers.csv"):
    mad_data = data.copy()
    sample_columns = data.columns[4::3]

    mad_data["cohort_median"] = mad_data[sample_columns].median(axis=1)
    print("[LOG] Calculated cohort median for each gene")

    for sample in sample_columns:
        mad_column_name = f"MAD_{sample}"
        mad_data[mad_column_name] = (mad_data[sample] -
mad_data["cohort_median"]).abs()
        print(f"[LOG] Calculated MAD values for {sample}")

    average_mad_values = {
```

```
        sample: mad_data[f"MAD_{sample}"].mean() for sample in sample_columns
    }

    output_columns = [f"MAD_{sample}" for sample in sample_columns] +
["cohort_median"]
    output_data = mad_data[output_columns]
    output_data.to_csv(file_name, index=False)
    print(f"[LOG] Saved MAD and cohort median data to '{file_name}'")

    return mad_data, average_mad_values


# Step 7-8: Apply FDR method to identify outliers and save results
def apply_fdr_method(average_mad_values, alpha=0.01):
    average_mad_df = pd.DataFrame(
        list(average_mad_values.items()), columns=["Sample", "Average MAD"]
    )

    rlm_model = RLM(
        average_mad_df["Average MAD"],
        np.ones_like(average_mad_df["Average MAD"]),
        M=HuberT(),
    )
    rlm_results = rlm_model.fit()

    residuals = average_mad_df["Average MAD"] - rlm_results.fittedvalues
    pvals = 2 * (1 - norm.cdf(residuals.abs() / rlm_results.scale))

    _, corrected_pvals, _, _ = multipletests(pvals, alpha=alpha, method="fdr_bh")
    average_mad_df["Outlier"] = corrected_pvals < alpha
    average_mad_df["Outlier"] = average_mad_df["Outlier"].apply(
        lambda x: "Yes" if x else "No"
    )

    print(f"[LOG] Applied FDR method with alpha={alpha}")
    return average_mad_df


def save_average_mad_with_outliers(average_mad_df,
file_name="step8_outliers.csv"):
    average_mad_df.to_csv(file_name, index=False)
    print(f"[LOG] Saved average MAD values with outlier status to '{file_name}'")
```

```python
# Step 9-14: Rescale cohorts to have a global median of 7, adjust log2 values,
and handle outliers
def rescale_cohorts(data, outlier_samples):
    sample_columns = data.columns[4::3]

    cohort_medians = data[sample_columns].apply(
        lambda row: np.median(
            [
                val
                for i, val in enumerate(row)
                if sample_columns[i] not in outlier_samples
            ]
        ),
        axis=1,
    )

    data["Cohort Median Excl. Outliers"] = cohort_medians
    filtered_medians = cohort_medians[cohort_medians > -6.6]
    global_median = np.median(filtered_medians)
    delta = 7 - global_median

    data["Adjusted Cohort Median"] = np.where(
        cohort_medians > -6.6, cohort_medians + delta, cohort_medians
    )
    return data, delta


def adjust_log2_fpkm_uq(data, delta, file_name="step14_adj_log2_fpkm_uq.csv"):
    adjusted_data = data.copy()
    sample_columns = data.columns[4::3]

    for column in sample_columns:
        adjusted_data[column] = np.where(
            adjusted_data[column] > -6.6,
            adjusted_data[column] + delta,
            adjusted_data[column],
        )
        print(f"[LOG] Applied delta adjustment to {column}")

    adjusted_data.to_csv(file_name, index=False)
    print(f"[LOG] Saved adjusted Log2 FPKM-UQ data to '{file_name}'")
    return adjusted_data


# Step 15: Adjust log2 values for all samples
```

```python
def adjust_all_samples(data, delta,
file_name="step15_final_adjusted_log2_fpkm_uq.csv"):
    final_adjusted_data = adjust_log2_fpkm_uq(data, delta)
    final_adjusted_data.to_csv(file_name, index=False)
    print(f"[LOG] Saved final adjusted Log2 FPKM-UQ data to '{file_name}'")
    return final_adjusted_data


# Step 15a: Replace negative log2 values with zero
def replace_negative_values_with_zero(
    data, file_name="step15a_replace_negative_with_zero.csv"
):
    adjusted_data = data.copy()
    sample_columns = data.columns[4::3]

    for column in sample_columns:
        adjusted_data[column] = adjusted_data[column].apply(lambda x: max(x, 0))
        print(f"[LOG] Replaced negative values with zero in {column}")

    adjusted_data.to_csv(file_name, index=False)
    print(f"[LOG] Saved data with negative values replaced with zero to
'{file_name}'")
    return adjusted_data


# Step 16: Validate re-scaling global median and document adjustments
def validate_rescaling(
    data, outlier_samples, initial_global_median,
file_name="rescaling_validation.txt"
):
    rescaled_data, new_delta = rescale_cohorts(data, outlier_samples)
    new_global_median = np.median(rescaled_data["Adjusted Cohort Median"])

    with open(file_name, "w") as f:
        f.write(f"Initial Global Median: {initial_global_median}\n")
        f.write(f"New Adjusted Global Median: {new_global_median}\n")

    print(f"[LOG] Documented global median rescaling in '{file_name}'")
    return rescaled_data, new_global_median


# Step 17: Save final output
def save_final_output(data, file_name="final_adjusted_log2_fpkm_uq.csv"):
    output_columns = ["gene_id", "Symbol"] + list(data.columns[4::3])
    final_output_data = data[output_columns]
```

```
    final_output_data.to_csv(file_name, index=False)
    print(f"[LOG] Saved final output to '{file_name}'")
    return final_output_data


# Step 18: Convert log2 values back to linear space
def convert_log2_to_linear(data, file_name="step18_log2_to_linear.csv"):
    linear_data = data.copy()
    sample_columns = data.columns[4::3]

    for column in sample_columns:
        linear_data[column] = (2 ** linear_data[column]) - 1
        print(f"[LOG] Converted log2 values back to linear space in {column}")

    linear_data.to_csv(file_name, index=False)
    print(f"[LOG] Saved linearized data to '{file_name}'")
    return linear_data


# Main function to run all steps
def main():
    combined_data_file = "combined_data.csv"
    gene_reference_file = "gene_reference.csv"
    pickle_file = "combined_data.pkl"

    combined_data = load_combined_data(combined_data_file)
    gene_reference = load_gene_reference(gene_reference_file)

    filtered_data = filter_protein_coding_genes(combined_data, gene_reference)
    percentiles, muq = calculate_75th_percentile(filtered_data)

    # Apply gene size transformation and calculate median gene sizes after the
transformation
    transformed_data, overall_mgs = apply_gene_size_transformation(filtered_data)

    # Normalize FPKM-UQ
    normalized_data = normalize_fpkm_uq(transformed_data, percentiles, muq,
overall_mgs)

    # Log2 transformation of FPKM-UQ
    log2_transformed_data = log2_transform_fpkm_uq(normalized_data)

    # Calculate MAD and identify outliers
    mad_data, average_mad_values = calculate_mad_and_identify_outliers(
        log2_transformed_data
```

```
    )

    # Apply the FDR method to identify outliers
    average_mad_df = apply_fdr_method(average_mad_values)

    # Save the average MAD values with outlier status
    save_average_mad_with_outliers(average_mad_df)

    # Rescale cohorts, adjust log2 values, and save the adjusted data
    rescaled_data, delta = rescale_cohorts(
        log2_transformed_data,
        average_mad_df[average_mad_df["Outlier"] == "Yes"]["Sample"].tolist(),
    )
    adjusted_data = adjust_all_samples(log2_transformed_data, delta)

    # Replace negative log2 values with zero
    data_with_zeros = replace_negative_values_with_zero(adjusted_data)

    # Validate rescaling and document the global medians
    final_rescaled_data, new_global_median = validate_rescaling(
        data_with_zeros,
        average_mad_df[average_mad_df["Outlier"] == "Yes"]["Sample"].tolist(),
        overall_mgs,
    )

    # Save the final output with adjusted Log2 FPKM-UQ values
    save_final_output(final_rescaled_data)

    # Convert log2 values back to linear space
    convert_log2_to_linear(data_with_zeros)

    # Clean up the Pickle file at the end to save space
    if os.path.exists(pickle_file):
        os.remove(pickle_file)
        print(f"[LOG] Deleted Pickle file: {pickle_file}")


if __name__ == "__main__":
    main()
```

**zeros_binomial.ipynb**

```
from scipy.stats import binom
import numpy as np
import pandas as pd
```

```
# Load the file to examine its structure
file_path = 'Zeros.csv'
data = pd.read_csv(file_path)

# Display the first few rows to understand its format
data.head()

# Define the function for Benjamini-Hochberg correction
def benjamini_hochberg(p_values, fdr_threshold=0.1):
    # Sort p-values and get their corresponding indices
    sorted_indices = np.argsort(p_values)
    sorted_p_values = np.array(p_values)[sorted_indices]

    # Number of hypotheses
    m = len(p_values)

    # Calculate the Benjamini-Hochberg thresholds
    bh_thresholds = np.array([(i + 1) / m * fdr_threshold for i in range(m)])

    # Initialize the adjusted p-values
    adjusted_p_values = np.zeros(m)

    # Calculate the adjusted p-values
    for i in range(m):
        adjusted_p_values[i] = sorted_p_values[i] * m / (i + 1)

    # Ensure adjusted p-values are non-decreasing
    adjusted_p_values = np.minimum.accumulate(adjusted_p_values[::-1])[::-1]

    # Ensure p-values don't exceed 1
    adjusted_p_values = np.minimum(adjusted_p_values, 1.0)

    # Return the p-values in their original order
    return adjusted_p_values[np.argsort(sorted_indices)]

# Calculate the binomial probability for each gene and get p-values
p_values = []

for index, row in data.iterrows():
    # Calculate the binomial probability of getting >= Observed_zeros successes
    observed_successes = row['Observed_zeros']
    total_samples = row['Total_samples']
    probability = row['Probability']
```

```
    # Calculate cumulative probability of getting at least observed_successes
    p_value = binom.sf(observed_successes - 1, total_samples, probability)
    p_values.append(p_value)

# Apply Benjamini-Hochberg correction
adjusted_p_values = benjamini_hochberg(p_values, fdr_threshold=0.1)

# Add raw and adjusted p-values to the dataframe
data['Raw_P_value'] = p_values
data['Adjusted_P_value'] = adjusted_p_values

# Save the updated dataframe to a CSV file
output_path = 'Zeros_with_P_values.csv'
data.to_csv(output_path, index=False)

# Display the updated dataframe
print(data)
```
