## Supplemental Figures for "Reliable RNA-seq analysis from FFPE specimens as a means to accelerate cancer-related health disparities research"

### Supplementary Figure 1

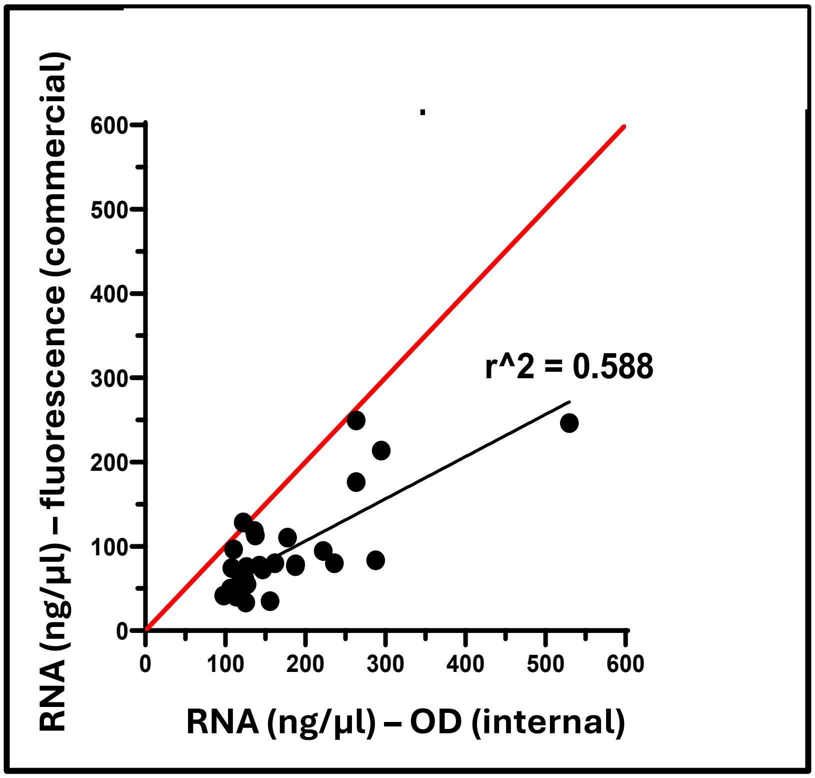

**Supplementary Figure 1. RNA quantitation.** RNA concentrations measured via fluorescence compared to internal laboratory measures using optical density (OD).

### Supplementary Figure 2

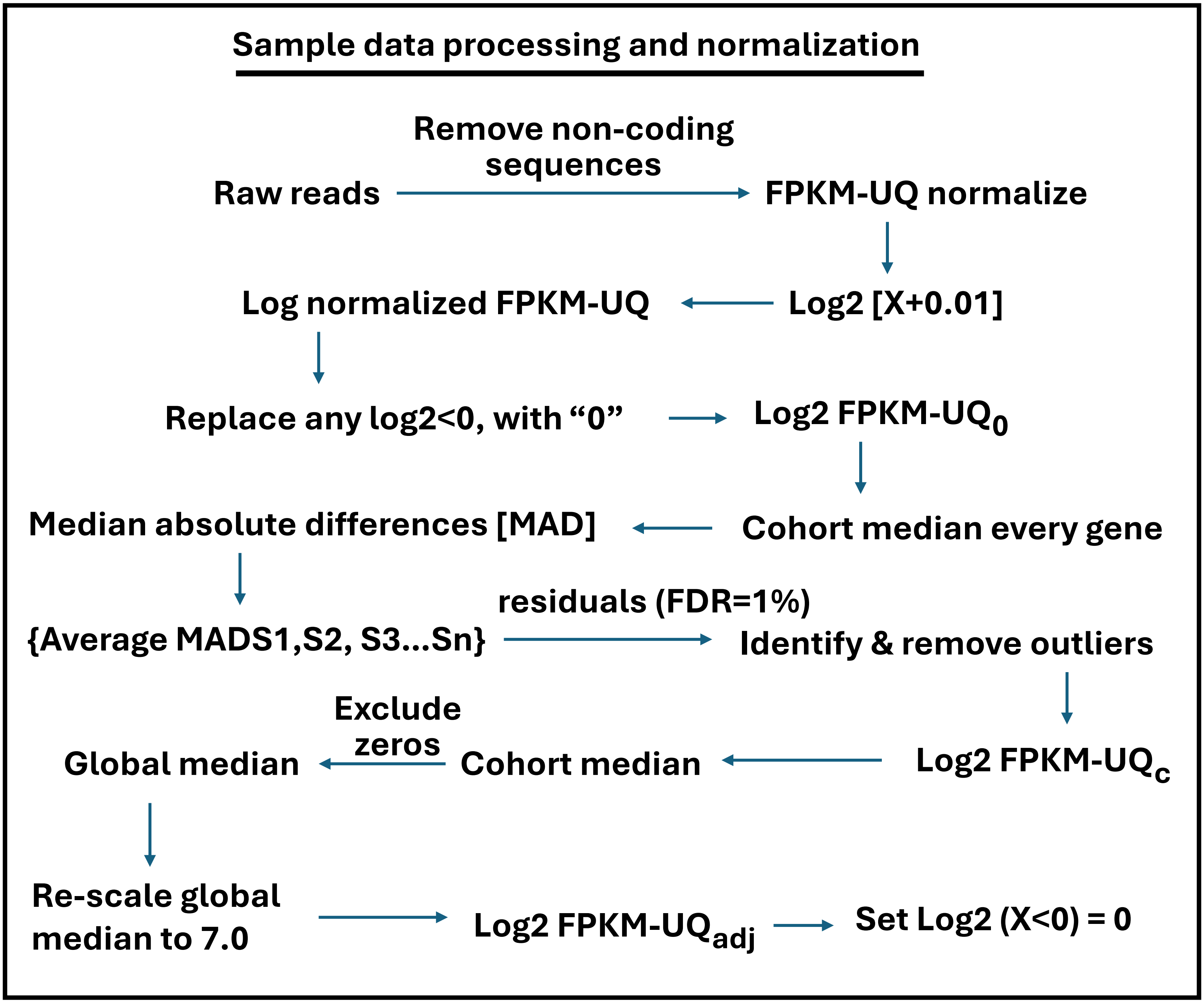

**Supplementary Figure 2. Analytical pipeline**. Pipeline for analysis of RNA-seq data inclusive of normalization.

### Supplementary Figure 3

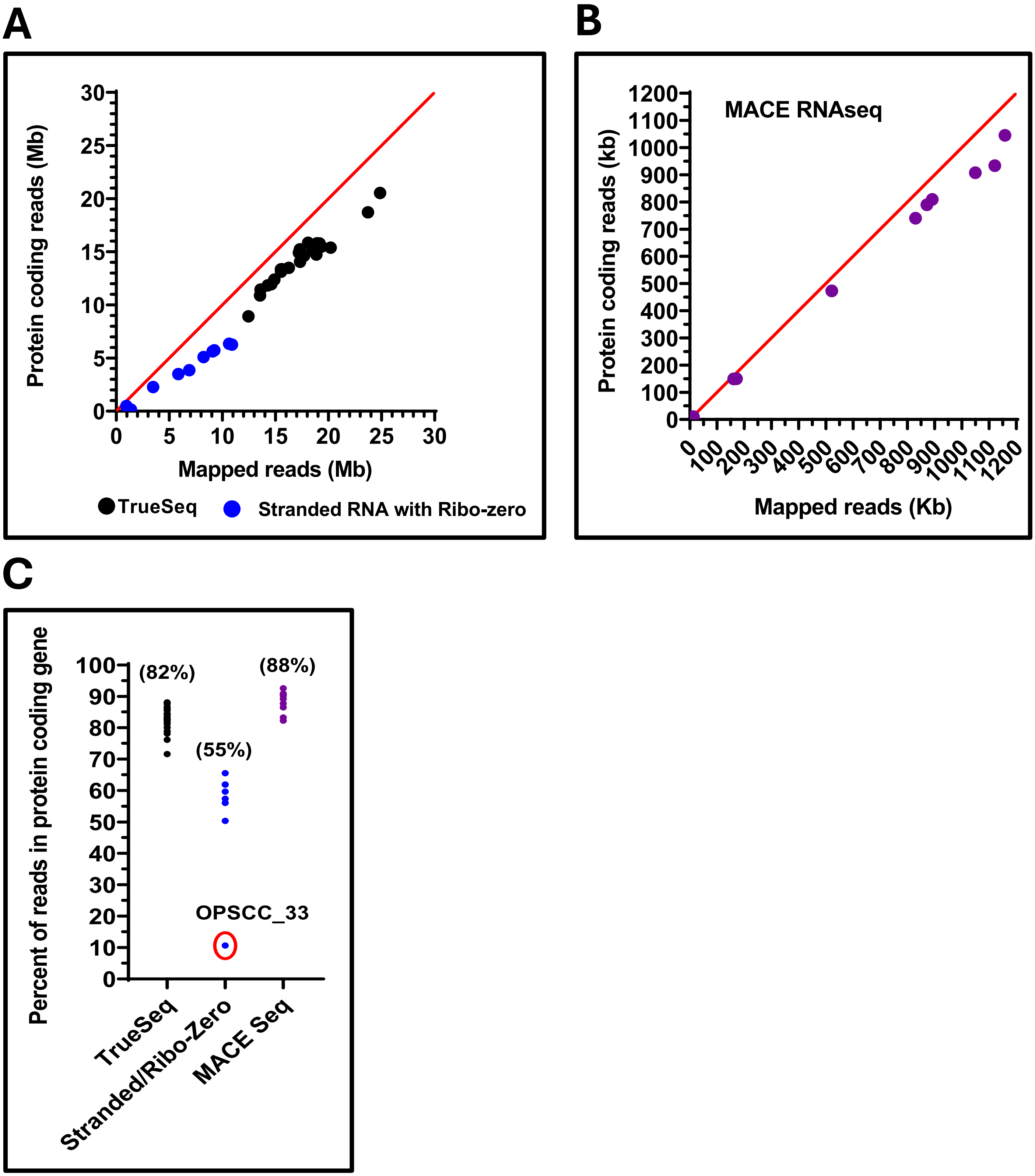

**Supplementary Figure 3. Coding vs non-coding reads**. Quantification of coding vs non-coding reads mapped using the True-Seq and Stranded RNA with Ribo-zero platforms (A) and the MACE RNAseq platform (B). Head to head comparison across platforms (C). Sample OPSCC_33, which had the lowest % of protein coding reads (red circle) was also a technical outlier (Supplementary Table 5)

### Supplementary Figure 4

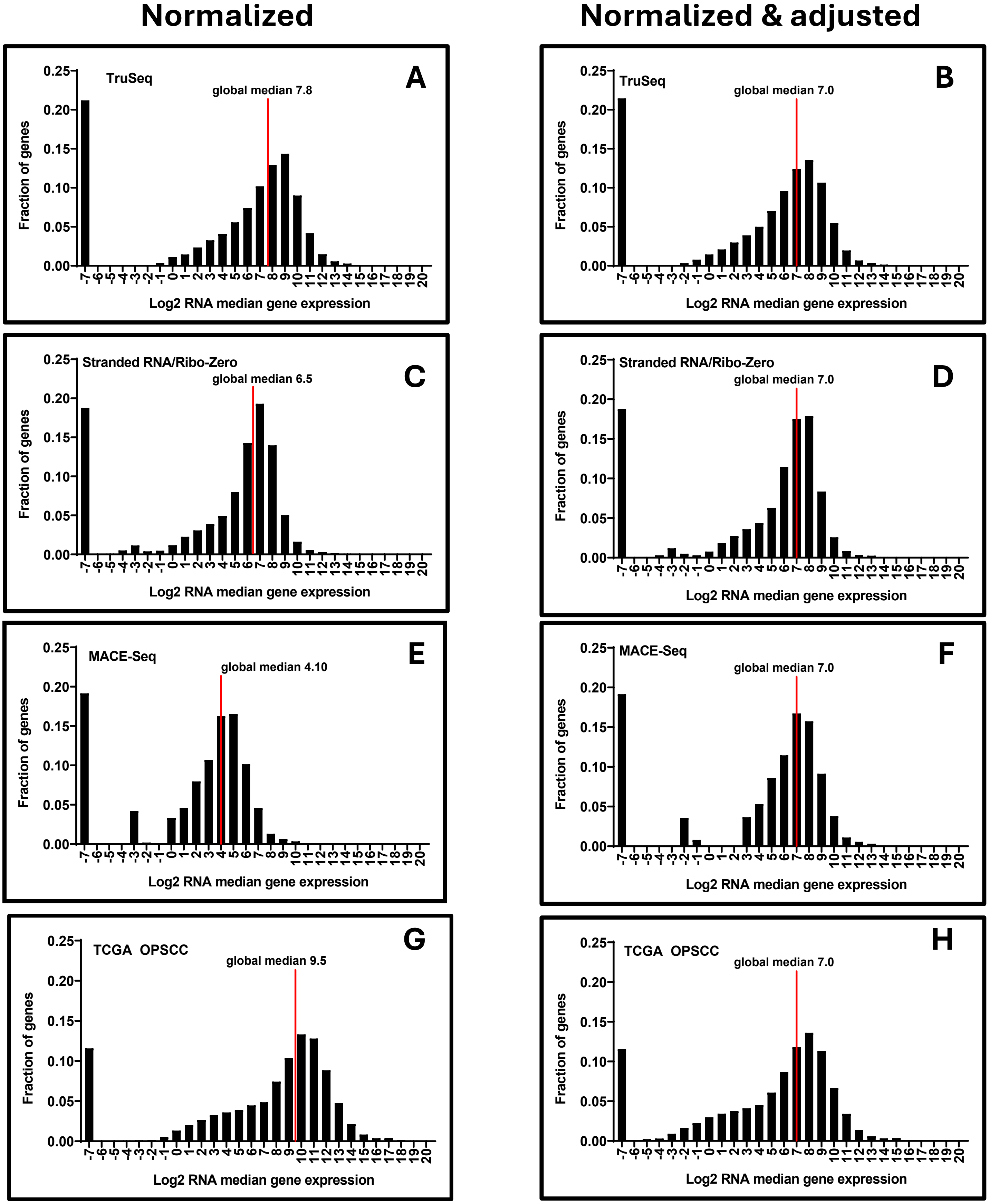

**Supplementary Figure 4. Distribution of RNA median gene expression**. The distributions of gene expression (i.e., median UQ normalized log 2 values) for each platform/cohort, along with the TCGA OPSCC RNA-Seq dataset similarly normalized **(A,C,E,G)** before global rescaling. Histograms for the rescaled cohorts were also generated **(B,D,F,H).**

### Supplementary Figure 5

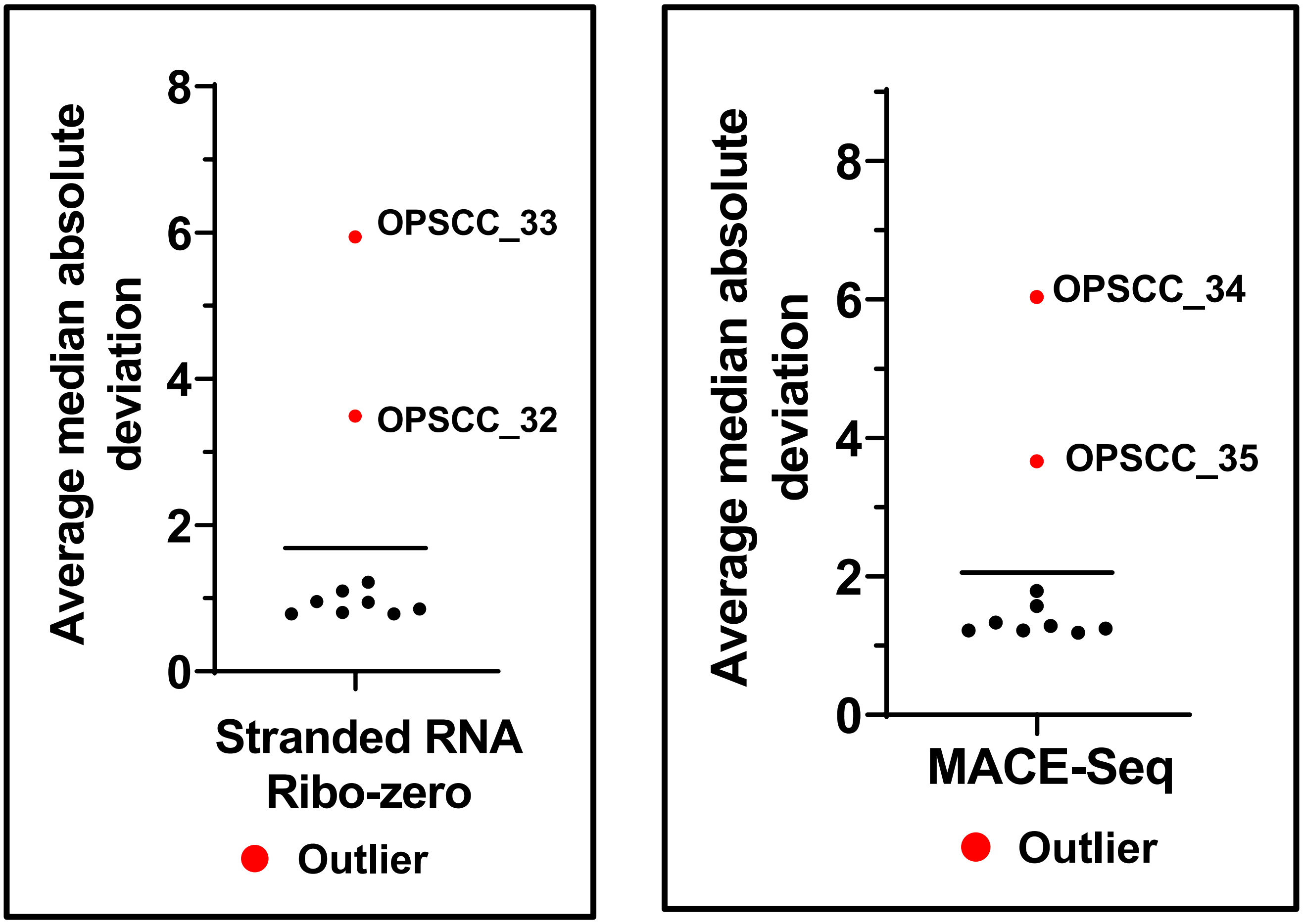

**Supplementary Figure 5. Average Median Absolute Deviation (MAD).** MAD values were averaged across all genes to calculate a sample specific average MAD value. Samples OPSCC_32 and 33 (Stranded RNA/Ribo-zero), OPSCC_34 and 35 (MACE-Seq) were identified as outliers with significantly different MAD values (Supplementary Table 5).

### Supplementary Figure 6

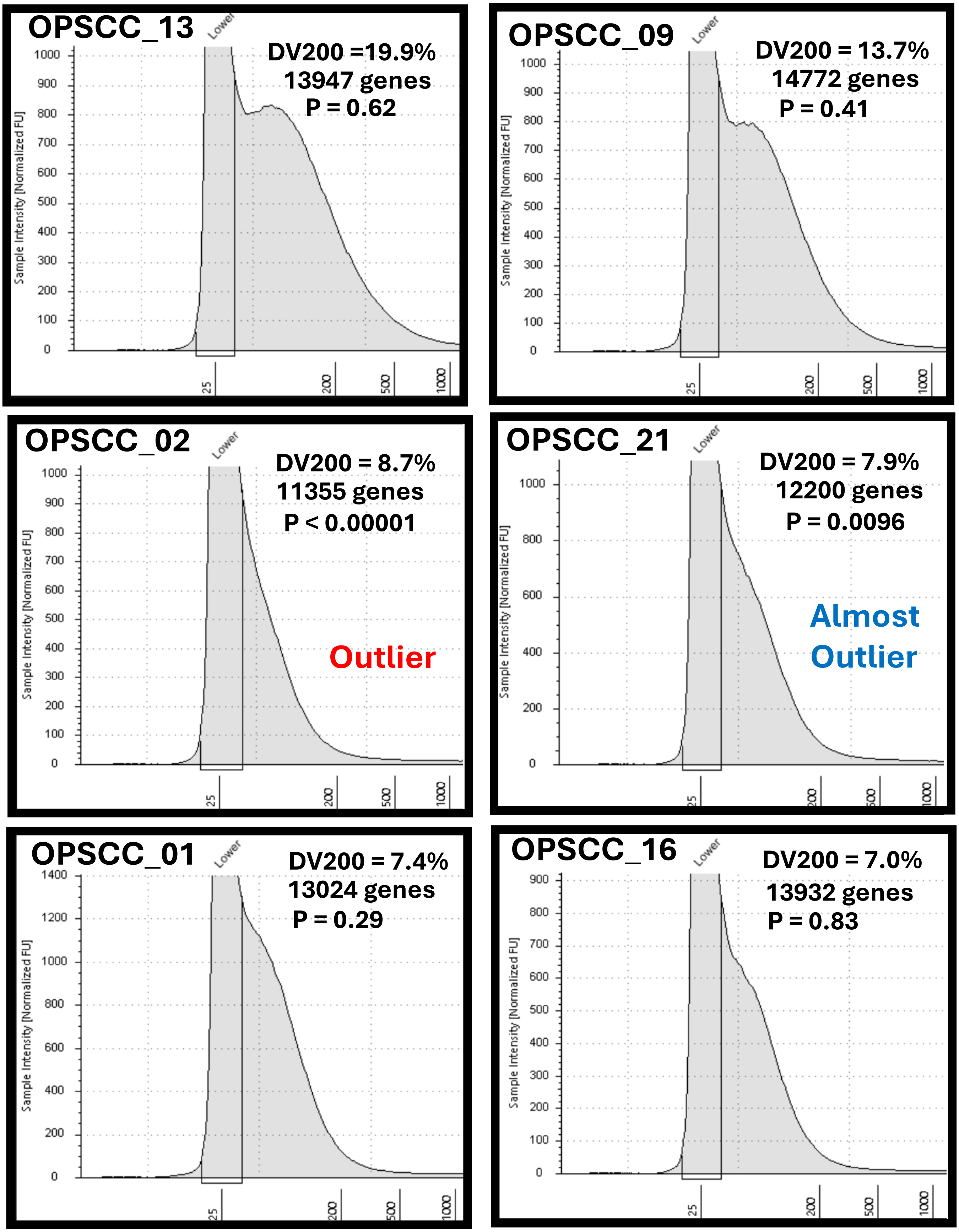

**Supplementary Figure 6. Relationship between DV200 and biological richness of gene expression**. DV200 values and RNA fragment size are shown for representative samples along with the relative number of usable genes and associated p-values from outlier analysis (Supplementary Table 5).

### Supplementary Figure 7

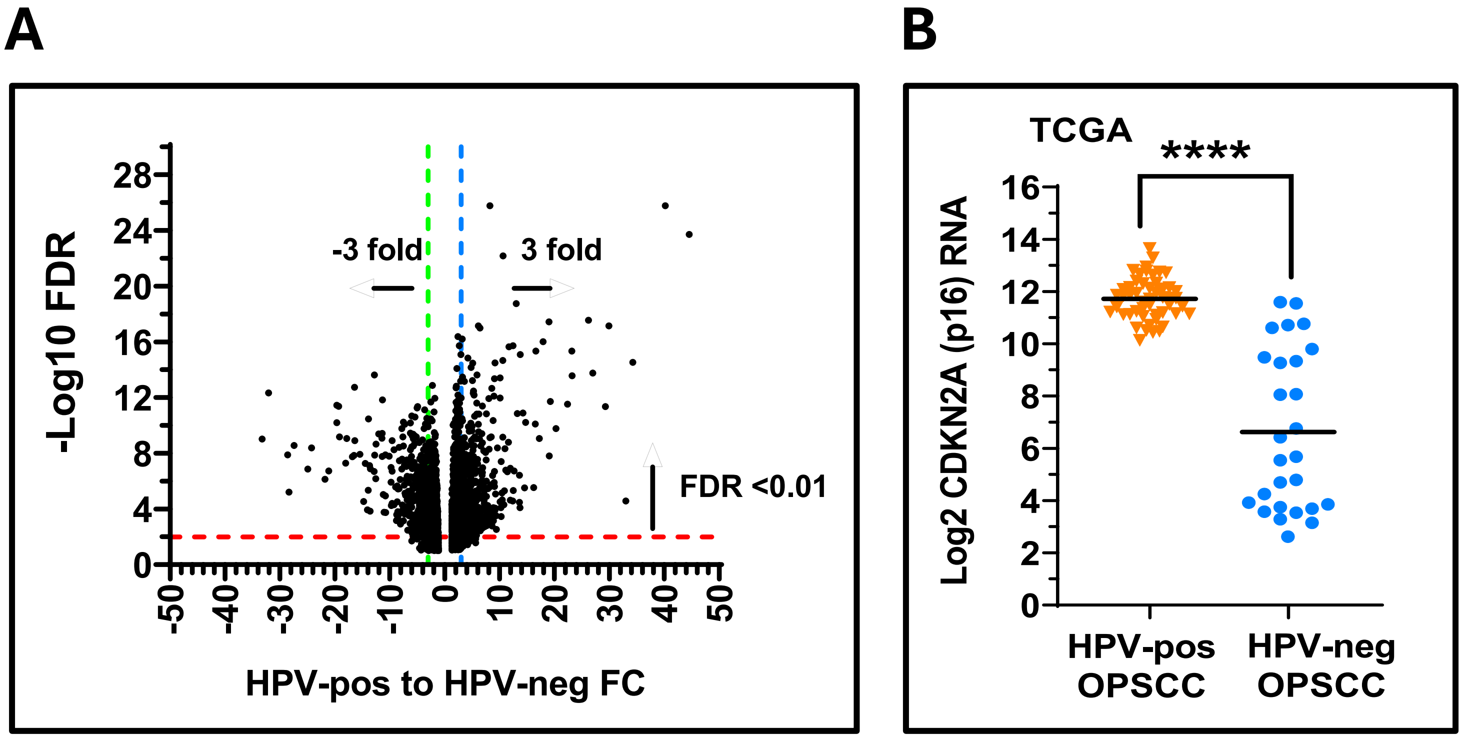

**Supplementary Figure 7. Correlation of HPV-associated genes and CDKN2A expression in TCGA samples**. A) Genes that showed ≥ 3-fold significant (FDR <0.1) difference up or downregulation based on HPV status in the OPSCC TCGA cohort were identified. B) Confirmation that CDKN2A expression is highly upregulated in HPV-associated (HPV-pos) TCGA samples. ****P< 0.0001.

### Supplementary Figure 8

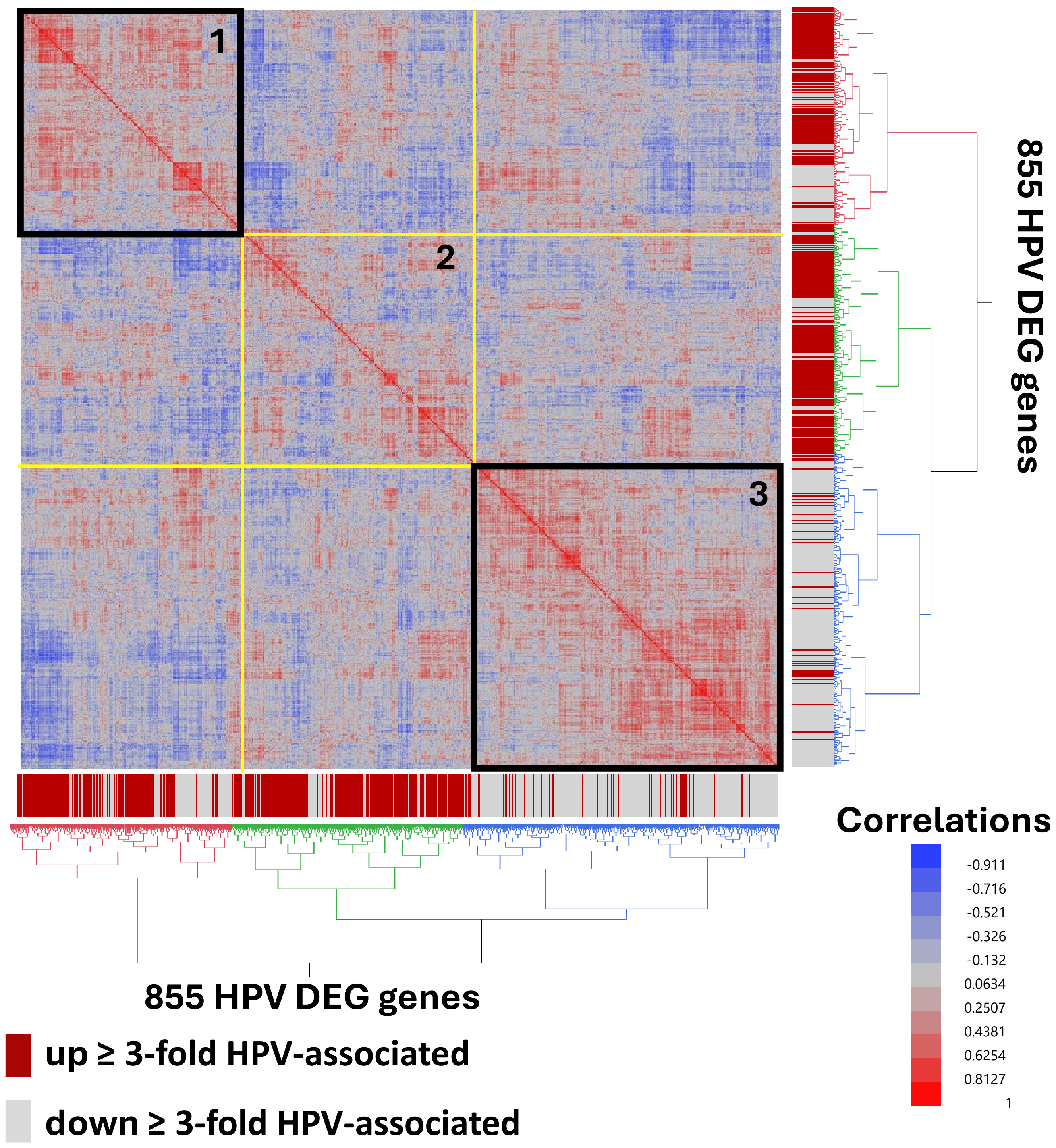

**Supplementary Figure 8. Cross-correlation of HPV associated genes**. Cross-correlation coefficients of gene expression values within the TruSeq cohort, using the list of 855 DEGs previously associated with HPV status in the TCGA OPSCC samples, were used for unsupervised clustering to Identify modules of genes (black boxes) that behaved similarly. Gene clusters 1 and 3 behaved robustly. Genes are annotated vertically and horizontally by whether they were upregulated (red boxes) or downregulated (grey boxes) in the original TCGA cohort according to HPV status.

### Supplementary Figure 9

A

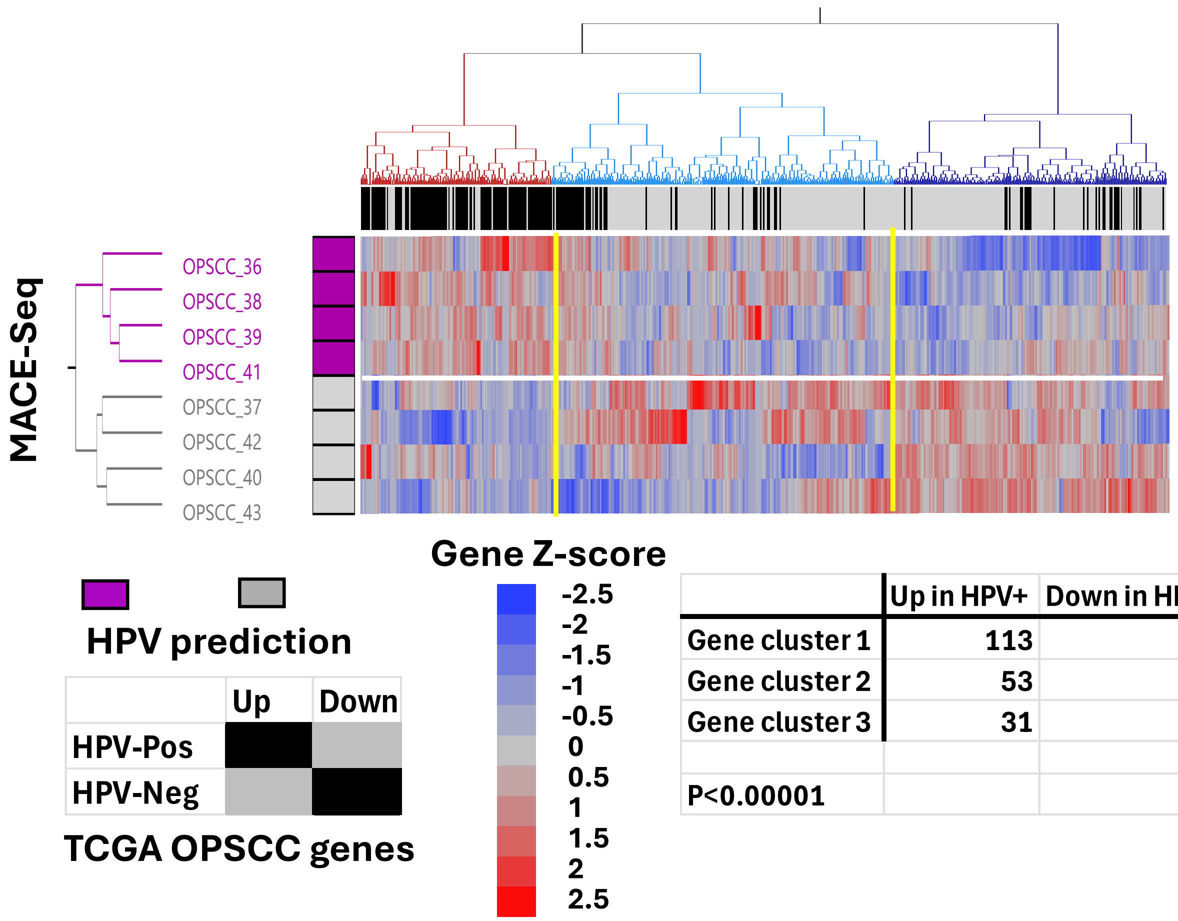

| | Up in HPV+ | Down in HPV+ |
|---|---|---|
| Gene cluster 1 | 113 | 23 |
| Gene cluster 2 | 53 | 192 |
| Gene cluster 3 | 31 | 164 |
| P<0.00001 | | |

B

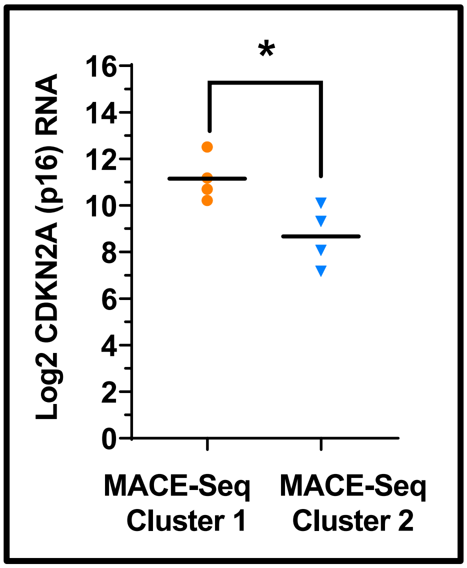

**Supplementary Figure 9. Correlation of HPV associated genes in the MACE-Seq cohort.** . Expression of HPV-associated DEGs (identified from the TCGA OPSCC reference cohort) was used for unsupervised 2-way clustering of MACE-Seq cohort samples (i.e., Ward's agglomerative hierarchical clustering) to predict HPV status (A). Samples predicted to be HPV-associated (i.e., HPV-Pos) are annotated with purple boxes. The regulation status of genes is annotated across the top of the heatmap with black boxes if they were also upregulated in HPV-associated TCGA samples or grey boxes if they were upregulated in HPV-independent (i.e., HPV-Neg) TCGA samples. Significant enrichment of genes upregulated in TCGA HPV-associated cancers was found in gene cluster 1 (red cluster) and enrichment of genes upregulated in HPV-independent TCGA samples is found in gene cluster 3 ( dark blue cluster), demonstrating significant separation (e.g., P <0.00001 by Chi-square testing). Specimens from sample cluster 1 (purple) predicted to be HPV-associated based on their gene expression pattern had higher CDKN2A (p16) expression than specimens from sample cluster 2 (grey) predicted to be HPV-independent (B). * P<0.05
